# Natriuretic peptides increase cGMP around cardiomyocyte mitochondria and protect against apoptosis

**DOI:** 10.1101/2022.08.22.504735

**Authors:** Gaia Calamera, Bernadin Ndongson-Dongmo, Dulasi Arunthavarajah, Mette Ovesen, Choel Kim, Finn Olav Levy, Kjetil Wessel Andressen, Lise Román Moltzau

## Abstract

Natriuretic peptides (NPs) increase cGMP, show beneficial cardiovascular effects and regulate energy metabolism in other tissues. However, little is known about their direct effect on cardiac mitochondria and cardiomyocyte apoptosis. Here, we examined whether NPs increase cGMP around mitochondria and alter apoptosis in cardiomyocytes. We constructed a novel FRET-based biosensor with high selectivity towards cGMP and found that ANP and CNP increase cGMP at the outer mitochondrial membrane. Moreover, ANP and CNP increased phosphorylation of the pro-apoptotic protein Drp1 and CNP prevented fragmentation of mitochondria. Stimulating cardiomyocytes with ANP or CNP reduced apoptosis, caspase 9 activation and cytochrome c release, suggesting that NPs decrease apoptosis through the intrinsic pathway that involves mitochondria. We suggest that cGMP increase in the outer mitochondrial membrane microdomain that inhibits the pro-apoptotic protein Drp1, leading to reduced mitochondrial fragmentation and thereby reduced apoptosis.

## 1. Introduction

Heart failure is associated with an imbalance in cell energy, changes in the energy source and oxidative stress^1–3^. In addition to providing ATP for the contractile machinery in cardiomyocytes, mitochondria are also important in regulation of cellular metabolism, cell differentiation, motility, cellular Ca^2+^ handling and apoptosis^4, 5^. The signaling molecule cyclic GMP (cGMP) has been shown to induce mitochondrial potassium channel opening, mediating a beneficial effect in ischemia-reperfusion injury^6–10^. Natriuretic peptides (NP) increase cGMP by activation of the particulate guanylyl cyclase (pGC) GC-A, activated by atrial (ANP) and brain NP (BNP), or GC-B, activated by C-type NP (CNP). NPs can mediate several beneficial effects in the cardiovascular system such as diuresis, natriuresis, vasodilation and antihypertrophic effects^11^. Consequently, recent attention has been given to drugs that increase NP levels as potential treatment of heart failure. Sacubitril-valsartan, a neprilysin (NEP) inhibitor in combination with an AT_1_ angiotensin II receptor antagonist, was shown to be superior to the angiotensin converting enzyme inhibitor enalapril in the treatment of heart failure with reduced ejection fraction (HFrEF)^12^. The role of NPs in heart disease still needs to be better understood to further improve treatment. Natriuretic peptides have been shown to regulate cell metabolism and be beneficial against metabolic disease^13^, but little is known about direct effects of NPs on mitochondrial function in the heart.

Signaling of the cyclic nucleotides (CN) cAMP and cGMP induces rapid changes in molecular networks where transient events occur in specific subcellular compartments. Therefore, the development of FRET-based biosensors for CNs has been instrumental in detecting CN dynamics in real-time in cardiomyocytes and in specific subcellular locations^14^. In cardiomyocytes at basal conditions (no receptor stimulation), levels of cGMP are 100 times lower than cAMP^15–17^. Thus, cGMP biosensors need to have a high affinity and a suitable dynamic range to visualize changes in cGMP levels. Nonetheless, a high selectivity vs. cAMP is also required considering the high cAMP concentrations often present. Currently, among the cGMP FRET biosensors, the red-cGES-DE5 biosensor is the most suitable for measurements in adult cardiomyocytes as it shows high affinity and high selectivity for cGMP^16, 18^. However, this biosensor has a low dynamic range^19^. We previously developed the Yellow *Pf*PKG biosensor, which displayed similar high affinity for cGMP as the red-cGES-DE5 and with a higher dynamic range. However, *Pf*PKG displayed low selectivity vs. cAMP^19^. Therefore, we wanted to develop a novel biosensor for cGMP with high selectivity to provide a better tool for real-time monitoring of cGMP in specific nanodomains such as the mitochondria. Further, our aim was to investigate whether NPs increase cGMP in nanodomains surrounding the mitochondria and whether NPs can modulate mitochondrial function in cardiomyocytes.

In the present study, we developed a FRET-based biosensor selective for cGMP and confined this to the outer mitochondrial membrane (OMM), which allowed us to detect local cGMP increases from both ANP and CNP in real-time in cardiomyocytes. We show that NPs can increase phosphorylation of Drp1 and CNP can prevent mitochondrial fragmentation. Moreover, CNP and ANP showed protective effect from apoptosis, confirmed by reduced poly (ADP-ribose) polymerase (PARP) cleavage, and caused reduced cytochrome c release from mitochondria and reduced caspase 9 activation, suggesting that the reduction of apoptosis involves the mitochondrial pathway.

## Results

### Development of selective FRET-based cGMP biosensors targeted to mitochondria

We have previously shown that cGMP increases in different cellular nanodomains through GC-A and GC-B stimulation, using both untargeted and localized FRET-based biosensors in adult cardiomyocytes^19, 20^. To investigate whether NPs increase cGMP at the mitochondria, we developed FRET-based cGMP biosensors that measure cGMP in the cytosol and at the outer mitochondrial membrane. We first constructed a potentially brighter variant of the previously published cGi-500 biosensor^21^. cGi-500 is based on the two tandem cyclic nucleotide binding (CNB) domains of cGMP-dependent protein kinase Iα, flanked between CFP and YFP fluorophores. We replaced the acceptor fluorophore with Venus and termed this new biosensor cGi-500V (Fig. 1a). Its affinity (EC_50_) for cGMP was 510±42 nM (Fig. 1a) with a maximal FRET change of 53.2±1.4% (Fig. 1d).

**Figure 1.**
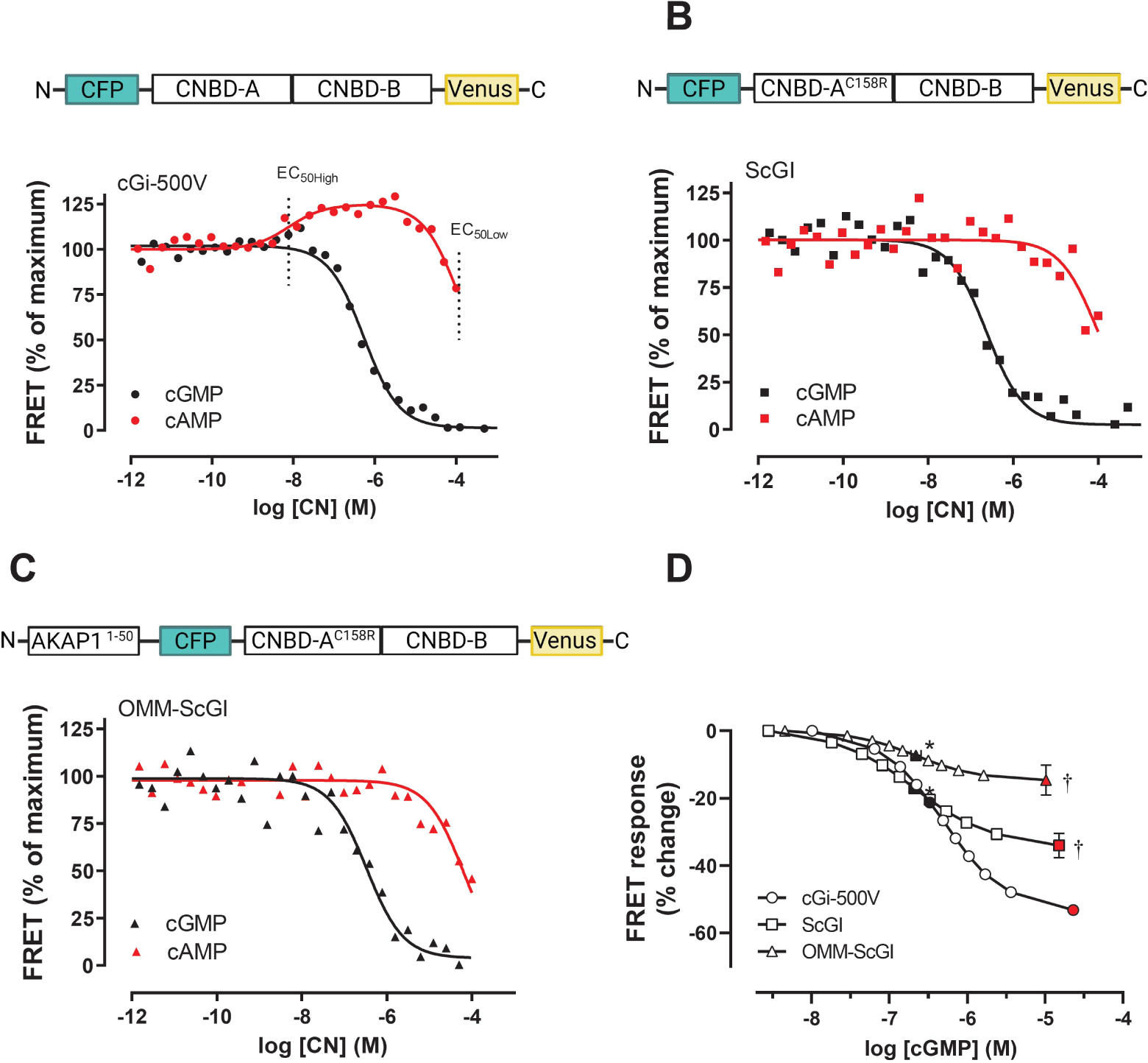
Development of FRET-based biosensor selective for cGMP and targeted to mitochondria. **A-C)** Upper panels: schematic of the indicated biosensor consisting of the fluorescent proteins CFP and Venus flanking the two cyclic nucleotide binding domains from PKG I without (cGi-500V) or with (ScGI) the C158R mutation. The OMM-ScGI contains an AKAP1 targeting sequence selective for the outer mitochondrial membrane. Lower panels: homogenates of HEK293 cells transfected with the indicated biosensor were incubated with increasing concentrations of either cGMP or cAMP. FRET (F_Venus_/F_CFP_) was measured as described in Materials and Methods and normalized to FRET in the absence of cGMP and at the highest cGMP concentration. Shown are representative of 10 (cGMP) and 5 (cAMP) independent experiments in **A**, 15 (cGMP) and 6 (cAMP) in **B**, 5 (cGMP) and 5 (cAMP) in **C. D)** Concentration-response curves generated by increasing concentrations of cGMP in homogenates of HEK293 cells expressing the indicated biosensors, as shown in a, b and c. EC_50_ values (marked as black filled symbols) and maximal change in FRET (marked as red filled symbols) are shown as mean±SEM from 12, 17 and 5 independent experiments for cGi-500V, ScGI and OMM-ScGI, respectively. *p<0.001 for *p*EC_50_ of ScGI or OMM-ScGI vs. cGI-500V; †p< 0.0001 for maximal FRET change of ScGI or OMM-ScGI vs. cGi-500V (one-way ANOVA, Sidak’s test).

Cyclic AMP is structurally similar to cGMP and some biosensors only display moderate selectivity between these nucleotides^19, 22^. Since cGMP can directly and indirectly modulate cellular concentrations of cAMP through inhibition or activation of certain PDEs^23, 24^, a biosensor with low cyclic nucleotide selectivity can indiscriminately bind either cyclic nucleotide and thus confound the results obtained. It is therefore essential to engineer a biosensor with high cGMP selectivity. First, we measured FRET of cGi-500V in the presence of increasing concentrations of cAMP and found that FRET *increased* by 23.7±2.4 % at low cAMP concentrations (EC_50High_ 12.1±1.8 nM), while FRET decreased at higher concentrations of cAMP (EC_50Low_ 106±28 µM) (Fig. 1a and Table 1) demonstrating that cAMP can bind cGi-500V and low levels of cAMP can thus interfere with the FRET responses for cGMP. The low selectivity of cGi-500V could reside in the CNB-A domain, that binds both cGMP and cAMP with similarly high affinities^25^. The CNB-B domain, on the other hand, is selective for cGMP and its cGMP-bound co-crystal structure showed that an arginine residue in β5 is responsible for this selectivity^26^. The corresponding position in CNB-A instead contains a cysteine^26^. We therefore mutated this cysteine (C158) in the CNB-A domain to an arginine and termed the mutant biosensor ScGI (<u>S</u>elective <u>c</u>yclic <u>G</u>MP kinase I). With ScGI, cAMP did not display a high affinity increase in FRET (as that obtained with the cGi-500V) and only a reduction of FRET was observed for the higher cAMP concentrations (EC_50_ of 125±23 µM; Fig. 3b and Table 1). The ScGI biosensor also displayed a ⁓2-fold *increased* affinity for cGMP (216±18 nM) compared to the cGi-500V resulting in ∼500-fold cGMP selectivity over cAMP. However, the mutation caused a reduction in the maximal FRET observed (34.0±3.6% *vs*. 53.2±1.4% for the cGi-500V, Fig. 1d).

**Table 1.** Sensitivity and dynamic range of the FRET biosensors. HEK293 cells expressing the indicated biosensor were homogenized and incubated with increasing concentration of cGMP or cAMP (where indicated) and fluorescence measured as described in Materials and Methods. The maximal FRET change was determined at the highest cGMP concentrations. EC_50_ values shown in µM were calculated from the average *p*EC_50_ values. Results shown are mean±SEM of 5-12 experiments. *p<0.001 and †p<0.0001 *vs*. cGi-500V.

| Sensor | cGMP |  | cAMP |  | cAMP |  | Max FRET change (%) |
| --- | --- | --- | --- | --- | --- | --- | --- |
| | $pEC_{50}$ | EC<sub>50</sub><br>(μM) | $pEC_{50High}$ | EC<sub>50</sub><br>(nM) | $pEC_{50Low}$ | EC<sub>50</sub><br>(μM) | |
| cGi-500V | 6.31±0.04 | 0.5 | 7.93±0.06 | 12 | 4.04±0.13 | 106 | 53.24±1.43 |
| ScGI | 6.68±0.03* | 0.2 | | | 3.97±0.03 | 125 | 34.08±3.62† |
| OMM-ScGI | 6.66±0.06* | 0.2 | | | 4.32±0.12 | 52 | 14.62±0.41† |

Since the ScGI with Cys158Arg no longer responded to the high affinity cAMP binding, and the affinity for cGMP closely matched previously published affinity for the isolated CNB-B^26^, we hypothesized that the cGMP-induced FRET change in both cGI-500V and ScGI was only due to binding of cGMP to the CNB-B. Since an arginine in β5 of CNB-B determines selectivity for cGMP^26^, we mutated this residue (R282A) in ScGI and observed a 10-fold reduction in cGMP affinity and a reduced dynamic range (19.6±1.2% change in FRET vs. 34% for ScGI) (Supplementary Fig. 1a). In addition, we determined if the presence of cAMP influenced cGMP-binding to the cGI-500V and ScGI biosensors in a competition-binding assay. In the presence of increasing concentrations of cGMP, adding a low concentration (1µM) of cAMP increased FRET of the cGI-500V of 25.0±1.7 %, but did not alter the affinity for cGMP (EC_50_ of 321±61 nM; Supplementary Table 1; Supplementary Fig. 1b). Adding a high concentration of cAMP (100µM) slightly reduced FRET (9.2±2.1 % reduction), and significantly decreased the potency of cGMP by ⁓2-fold (EC_50_ 1173±202 nM) (Supplementary Fig. 1b). For the ScGI biosensor, 1 µM cAMP did not affect the FRET response (7.3±3.6% increase, p=0.08) and did not modify the potency of cGMP (EC_50_ 294±116 nM). Adding 100µM cAMP significantly reduced FRET (37.2±7.3 % of maximum reduction) and reduced the cGMP potency by ⁓3-fold (EC_50_ 683±174 nM) (Supplementary Fig. 1c). Taken together, these data suggest that cAMP at low concentrations binds to the CNB-A of cGi-500V, which is not responsible for determining the cGMP affinity but rather induces a conformational change of the biosensor and therefore increases FRET. Cyclic AMP at high concentrations, on the other hand, competes with cGMP for binding to the CNB-B in both cGi-500V and ScGI. In addition, this confirms that the mutation in the CNB-A of the ScGI prevents the high affinity binding of cAMP displayed by the cGi-500V.

Next, we constructed a mitochondria-localized ScGI biosensor using the N-terminal targeting sequence of AKAP1, an anchoring protein known to be localized at the OMM and directing PKA signaling^27–29^. This biosensor was termed OMM-ScGI and displayed an affinity for cGMP similar to the untargeted ScGI biosensor (EC_50_ 225.7±32.3 nM) with ⁓200-fold selectivity *vs*. cAMP (EC_50_ 52±11 µM) (Fig. 1c and Table 1). However, adding this tag to the ScGI biosensor decreased the maximal change in FRET significantly compared to the untargeted sensor (14.6±0.4%, Fig. 1d).

### Real time measurement of cGMP around mitochondria

To determine if the OMM-ScGI biosensor displayed mitochondrial targeting, we expressed the ScGI and OMM-ScGI in H9c2 cells and in adult rat ventricular cardiomyocytes and determined if these co-stained with the mitochondrial marker Mitotracker Deep Red. The OMM-ScGI biosensor co-localized with Mitotracker Deep Red while the untargeted biosensor did not (Fig. 2a; Supplementary Fig.2a), suggesting that the OMM-ScGI localizes to the mitochondria.

**Figure 2.**
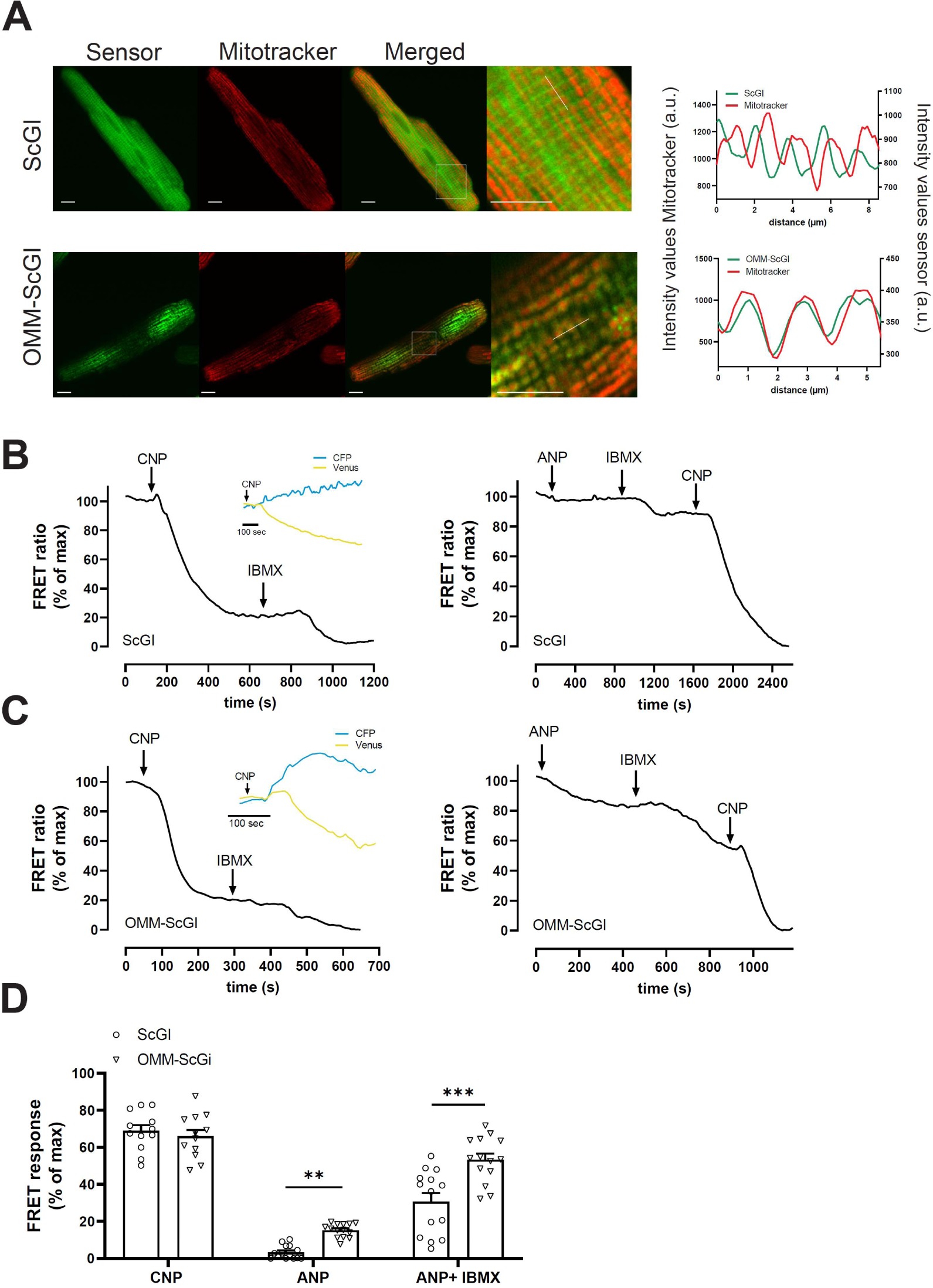
CNP and ANP increase cGMP at the outer mitochondrial membrane in cardiomyocytes. **A)** Cells expressing the ScGI (upper panel) and OMM-ScGI (lower panel), were stained with Mitotracker Deep Red, washed and visualized under a confocal microscope as described in Materials and Methods. Scale bar: 10µm. Right panels show colocalization by intensity overlay of both fluorescent signals (Mitotracker and sensors) from indicated dotted bar. a.u.: Arbitrary units **B and C)** Representative FRET ratio (Venus/CFP) recordings in single cells expressing the indicated biosensor and stimulated with CNP (300nM) and IBMX (100µM) (left panels) or ANP (1µM) followed by maximal activation of the biosensor (IBMX 100µM + CNP 300 nM) (right panels). The inserts in the left panels show corresponding traces of separate Venus and CFP intensities upon stimulation with CNP (300 nM). Shown are traces representative of 10-14 cells from 5-8 independent experiments **D)** Quantification of FRET responses from stimulation with CNP, ANP or combined response of IBMX + ANP. **p<0.01 and ***p<0.001 (one-way ANOVA, Sidak’s test).

Others have shown that stimulation of soluble guanylyl cyclase (sGC) increases cGMP at the OMM in neonatal cardiomyocytes using an OMM-localized FRET-based biosensor^30^, whereas it is unknown whether NPs elicit cGMP at the OMM. To assess if NPs increased cGMP around this localized biosensor, we stimulated single cells expressing either the ScGI or the OMM-ScGI biosensors with CNP (300 nM) followed by the non-selective PDE inhibitor IBMX (100 µM) to obtain maximal biosensor activation and measured FRET, reflecting cGMP levels (Fig. 2b and d; Supplementary Fig.2b and d). Stimulating ScGI-expressing cells with CNP decreased FRET (increase in cGMP) by 90.5±2.0% of maximum in H9c2 and by 68.8±3.1% of maximum in cardiomyocytes, while in OMM-ScGI expressing cells, CNP decreased FRET by 81.1±4.3% of maximum in H9c2 and by 66.6±4.3% of maximum in cardiomyocytes (Fig. 2b-d; Supplementary Fig. 2b-d). This suggests that GC-B stimulation causes an increase in cGMP in the cytosol, as previously shown^16, 19, 20^ and at the OMM; moreover IBMX gave a further increase in cGMP in both the cytosol and at the OMM, indicating that PDEs regulate these pools of cGMP (Fig. 2d; Supplementary Fig. 2d).

We and others have previously shown that GC-A and GC-B stimulation increases cGMP with different magnitudes and in different subcellular compartments^16, 19, 20, 31^. We therefore also wanted to measure the ANP-stimulated cGMP pool. Stimulation of ScGI-expressing cells with ANP (1µM) gave a FRET response of 2.4±1.5% of maximum in H9c2 and of 2.2±1.3% of maximum in cardiomyocytes (Fig. 2b and Supplementary Fig.2b). Further stimulation with IBMX gave a FRET response of 5.1±1.1% change in H9c2 and 29.2±3.8% change in cardiomyocytes (Fig. 2d and Supplementary Fig.2d). Stimulation of OMM-ScGI-expressing cells with ANP generated a larger response compared to the untargeted sensor, with 21.1±6.7 % of maximum in H9c2 and 15.2±1% of maximum in cardiomyocytes (Fig. 2c and Supplementary Fig.2c). IBMX gave a further FRET response of 16.1± 2.4% change in H9c2 and 39.1±2.9% change in cardiomyocytes (Fig. 2d and Supplementary Fig.2d). Both biosensors displayed an ANP-induced cGMP increase significantly lower than that induced by CNP, but ANP at the OMM gave a significantly larger response than in the cytosol in adult cardiomyocytes. These data taken together suggest that CNP induces a greater cGMP pool both in the cytosol and at the OMM, while ANP induces a smaller cGMP pool, which instead is restricted to the OMM in adult cardiomyocytes.

### NPs modulate Drp1 phosphorylation and prevent mitochondrial fragmentation

Mitochondrial function is modulated by phosphorylation/dephosphorylation of several proteins involved in mitochondrial dynamics. Among these, Drp1 is localized at the OMM and PKA-mediated phosphorylation of Drp1 at Ser637 was found to protect against apoptosis and mitochondrial fragmentation^28, 32–34^. We therefore investigated whether NPs modulate phosphorylation of Drp1 in adult cardiomyocytes and found that both CNP and ANP increased phosphorylation at Ser637, with CNP giving a significantly greater phosphorylation compared to ANP (Fig. 3a) without affecting the total protein expression (Supplementary Fig. 3).We next tested whether NPs can prevent mitochondrial fragmentation in H9c2 cells undergoing H_2_O_2_-mediated apoptosis. First, we measured CNP- and ANP-mediated cGMP increase in H9c2 cells over 24h and found that cGMP levels reached a peak already at 10 min and with an increase above basal maintained throughout 12h for both CNP and ANP (Supplementary Fig. 4). Therefore, we stimulated H9c2 cells with H_2_O_2_ (100µM) for 3h and found that mitochondria were significantly more fragmented compared to control cells (Fig. 3b) and that concomitant CNP-stimulation for 3h significantly inhibited mitochondrial fragmentation induced by H_2_O_2_ (Fig. 3b). ANP, on the other hand, did not modify H_2_O_2_-induced mitochondrial fragmentation (Fig. 3b), even though the proportion of cells with elongated mitochondria was slightly increased (30.4% with H_2_O_2_ + ANP *vs* 18.5% with H_2_O_2_; Table 2). All together, these data suggest that NPs modify mitochondrial dynamics.

**Figure 3.**
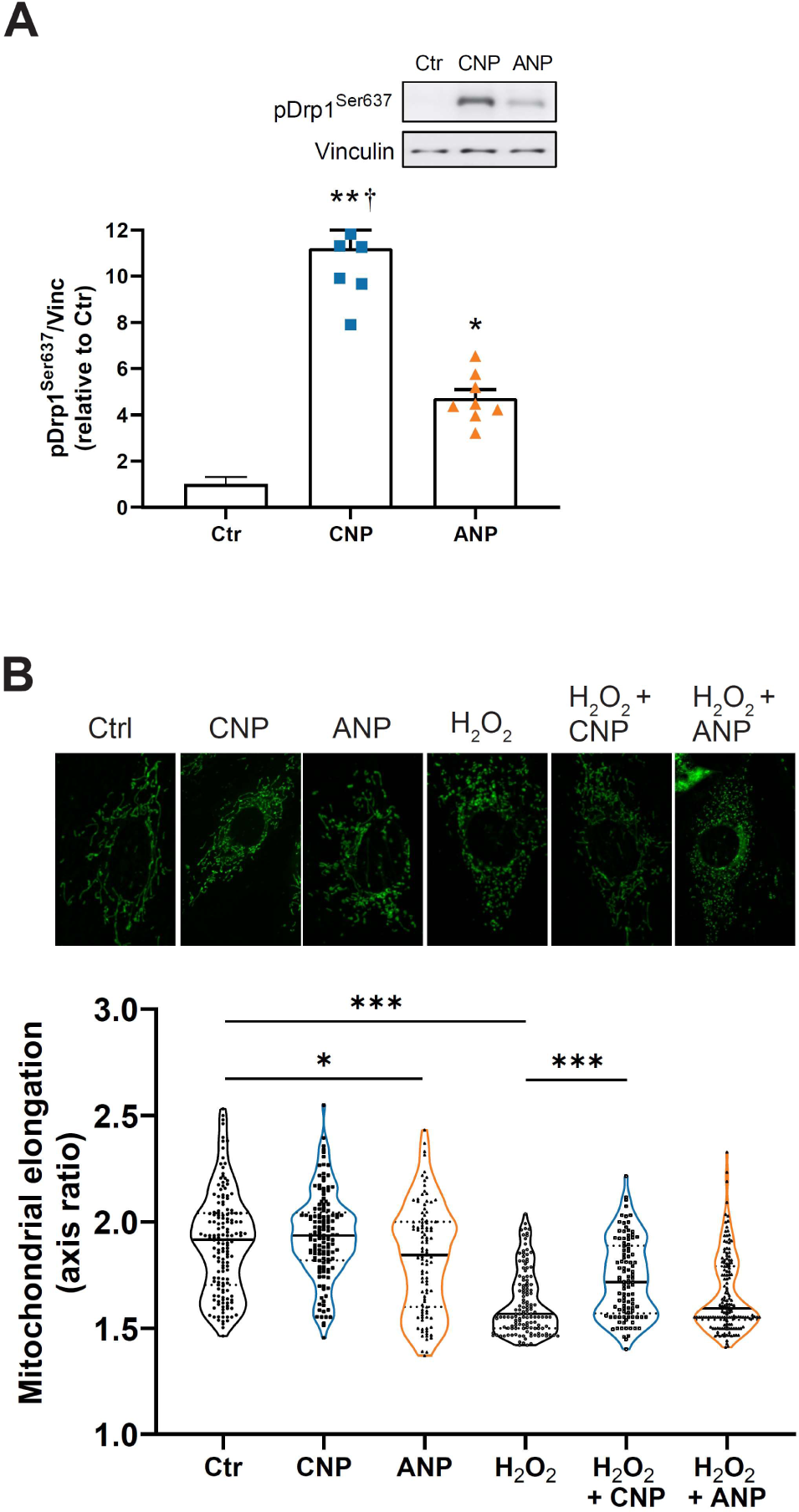
NPs modulate Drp1 phosphorylation and prevent mitochondrial fragmentation. **A)** Adult rat cardiomyocytes were stimulated with vehicle (Ctr), CNP (1 µM) or ANP (1 µM) for 3 h and lysed. Phosphorylation of Drp1 at Ser637 (pDrp1^Ser637^) was determined as described in Materials and Methods. The insert shows representative western blots of 5 independent experiments. The bands were quantified and normalized to loading control (vinculin) and Ctr (mean±SEM). *p< 0.05 and **p<0.01 *vs*. Ctr; †p<0.01 CNP *vs*. ANP (one-way ANOVA with Tukey’s test). **B)** H9c2 cells were stained with Mitotracker Green, washed and visualized under a confocal microscope and pictures processed by ImageJ to obtain the mitochondrial elongation as described in Materials and Methods. Data are shown as mean±SEM of 102-158 cells from 4 independent experiments. *p<0.05, ***p<0.001 (one-way ANOVA, Sidak’s multiple comparisons test).

**Table 2.** Fraction of fragmented *vs.* elongated mitochondria. Data from Figure 5 were distributed in two different fractions per each group, where cells with mitochondrial elongation (axis ratio) smaller than 1.75 were considered as having fragmented mitochondria and cells with mitochondria larger than 1.75 were considered as having elongated mitochondria. The threshold was chosen based on mitochondria size distribution in the Ctr group. Fractions are indicated as per cent of total number of cells per each group.

| **Mitochondrial elongation** | **Ctr** | **CNP** | **ANP** | **H<sub>2</sub>O<sub>2</sub>** | **H<sub>2</sub>O<sub>2</sub>+CNP** | **H<sub>2</sub>O<sub>2</sub>+ANP** |
| --- | --- | --- | --- | --- | --- | --- |
| <1.75<br>(fragmented) | 30.9% | 16.8% | 38.8% | 81.5% | 56.9% | 69.6% |
| >1.75<br>(elongated) | 69.1% | 83.2% | 61.2% | 18.5% | 43.1% | 30.4% |

### Natriuretic peptides increase cell viability

In order to further investigate the functional role of cGMP signaling at the OMM, we decided to study the effects of NPs on cell survival which is tightly connected to mitochondrial function. We first measured cell viability of cardiomyocytes cultured for 20 h to induce stress to the cells. During cell death, LDH is released to the medium, and this was measured over the next 24 h and progressively showed a time-dependent increase in cell death (Fig. 4a). To determine if NPs could modify cell viability, we stimulated cardiomyocytes with CNP (1µM) or ANP (1µM) and found that both natriuretic peptides significantly reduced LDH levels in the medium compared to vehicle over time (Fig. 4a). This indicates a protective effect of CNP and ANP against cardiomyocyte death. Further, we assessed the production of ATP as an indicator of metabolically active cells. ATP levels were increased by stimulation with CNP or ANP for 3h (1.6±0.1 and 1.8±0.2 fold increase *vs.* control, respectively) and 24h (1.4±0.1 and ANP:1.4±0.1 fold *vs.* control, respectively; Fig. 4b) suggesting that NPs increase cell viability, in line with our results on LDH measurements. To determine whether NPs increase cGMP over such prolonged periods (3-24h), cardiomyocytes were stimulated with CNP or ANP for up to 24 h and we found that CNP-stimulation increased cGMP compared to control over the entire 24 h showing peak cGMP levels around 1 h, while ANP-stimulation increased cGMP to a lesser extent than CNP with a peak around 30 min and decreased almost down to control levels between 12 and 24 h (Supplementary Fig. 5). These data suggest that cGMP increases are sustained and can mediate downstream effects on cell viability during prolonged NP stimulation.

**Figure 4.**
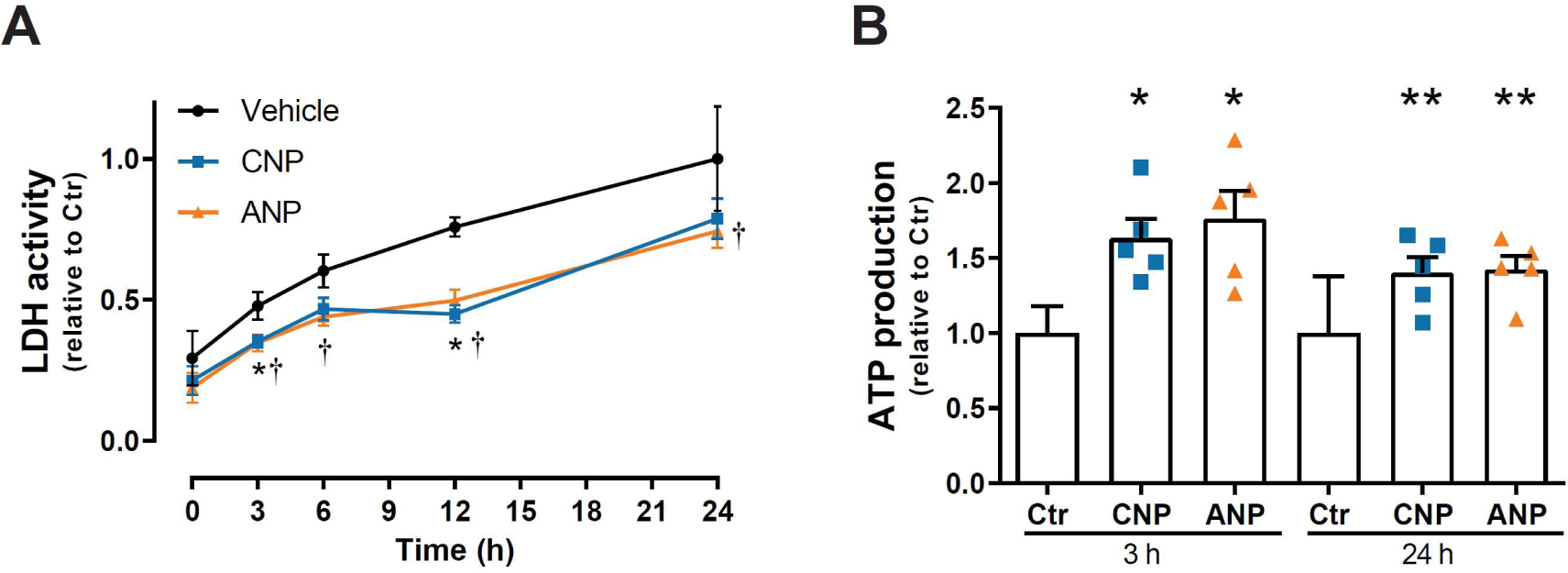
NPs reduce cell death and increase ATP levels compared to the control. **A)** Cardiomyocytes were stimulated with Vehicle, CNP (1µM) or ANP (1µM) for 24 h and sampling of the medium performed at the indicated time. LDH activity in the medium was determined as described in Materials and Methods. LDH activity at 24 h from Vehicle was used as control. Data shown are mean±SEM of 8 independent experiments. *(CNP) and † (ANP); p <0.05 compared to vehicle (two-way ANOVA with Dunnett’s multiple comparisons test). **B)** Cardiomyocytes were stimulated with vehicle (Ctr, CNP (1µM) or ANP (1µM) for the indicated times and ATP levels measured as described in Materials and Methods and normalized to Ctr (44±14×10^3^ RLU/µg protein at 3h and 26±10×10^3^ RLU/µg protein at 24h). Data shown are mean±SEM of 5 independent experiments. *p<0.05 and **p<0.01 compared to control (one-way ANOVA with Sidak’s test).

### NPs protect against apoptosis through the mitochondrial intrinsic pathway

LDH release from damaged cells could result from either necrosis or apoptosis. We therefore explored if NPs could alter apoptosis and performed TUNEL assay on adult cardiomyocytes. Stimulating cardiomyocytes for 3 h with either CNP or ANP reduced the number of apoptotic cells (76.1±5.3% and 65.9±8.3% compared to control, respectively; Fig. 5a). To verify the effects of NPs on apoptosis, we measured the cleavage of PARP, which will prevent PARP from repairing DNA damage and therefore induce apoptosis. Stimulation with CNP or ANP for 3 h significantly decreased PARP cleavage (Fig. 5b). Taken together, these data suggest that both CNP and ANP reduce apoptotic cell death.

**Figure 5.**
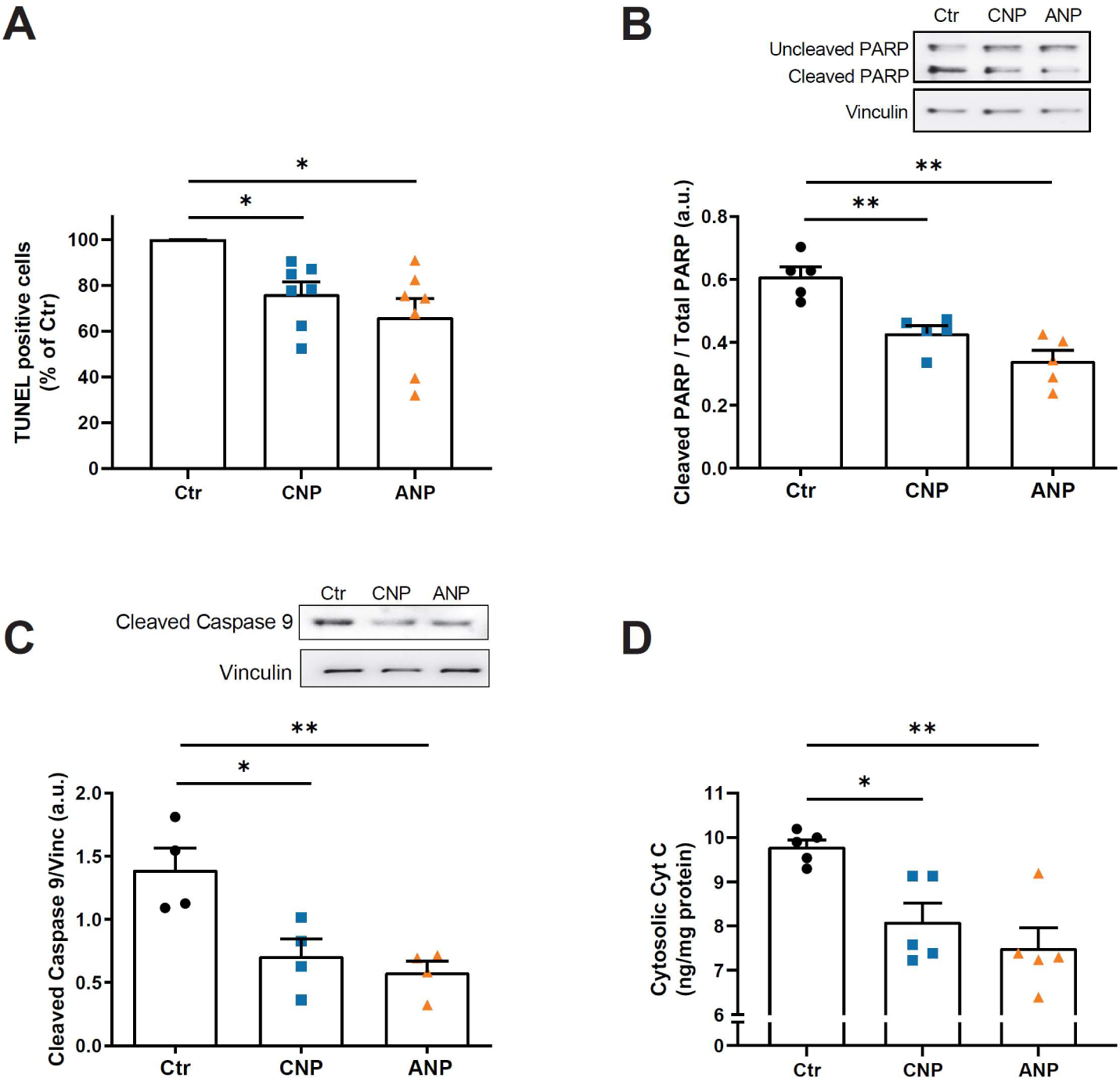
NPs protect against apoptosis through the mitochondrial intrinsic pathway. Cardiomyocytes were incubated with vehicle (Ctr), CNP (1µM) or ANP (1µM) for 3 h. **A)** Cells were fixed in 1% paraformaldehyde and fragmented DNA was stained by anti-digoxigenin antibody conjugated with fluorescein. Total number of cells and TUNEL positive cells were counted and shown as a per cent of vehicle treated cells (29.4 % ±3.2). Data shown are mean±SEM of 7 independent experiments. **B)** Cells were lysed and the amount of uncleaved and cleaved PARP and vinculin (loading control) were determined as described in Materials and Methods. Insert shows a representative blot. Data are mean±SEM of 5 independent experiments. **C)** Cells were lysed and active caspase 9 was determined as described in Materials and Methods and normalized to vinculin. Insert shows a representative blot. Quantification of cleaved caspase 9 as a fraction of the loading control (vinculin) and are shown as mean±SEM of 4 independent experiments. **D)** Cytochrome c levels were measured in the cytosolic fraction as described in Material and Methods. Values shown are mean±SEM of 5 independent experiments. *p<0.05 and **p<0.01 *vs*. Ctr (one-way ANOVA with Sidak’s (**A**) or Dunnett’s multiple comparisons (**B-D**) test. a.u.: Arbitrary units.

To investigate if the NP-mediated protective effect against apoptosis was through the intrinsic apoptotic pathway, we measured the activation of caspase 9, which depends on cytochrome c release from the mitochondria. CNP- and ANP-stimulation for 3 h significantly reduced the levels of cleaved caspase 9 compared to control (Fig. 5c). In addition, CNP and ANP significantly reduced levels of cytochrome c in the cytosol compared to the control (82.7% ±5.3 (CNP) and 76.7% ± 8.3 (ANP) of control, Figure 5d). Together, these data show that both NPs prevent cardiomyocyte apoptosis through the intrinsic mitochondrial pathway.

## Discussion

In this study, we developed a selective FRET-based biosensor for real-time detection of cGMP in intact cells and we targeted this to the OMM. We used this biosensor to show that ANP and CNP increase cGMP at the outer mitochondrial membrane in cardiomyocytes. In addition, both CNP and ANP increase Drp1 phosphorylation, with CNP also preventing mitochondrial fragmentation. Further, we show that the natriuretic peptides CNP and ANP protect against apoptosis of cardiomyocytes, with reduced PARP cleavage and increased cell viability. Moreover, we show that the natriuretic peptides reduce both activation of caspase 9 and release of cytochrome c from mitochondria to the cytosol, suggesting that the protective effects involve the mitochondrial intrinsic pathway.

### FRET-based biosensors detect cGMP increase at the mitochondrial outer membrane

Several FRET-based biosensors have been developed with different fluorescent proteins, cGMP binding domains and affinity/selectivity profiles. Among the most sensitive biosensors, we find the red-cGES-DE5, Yellow *Pf*PKG and Red *Pf*PKG biosensors^16, 19^, with cGMP affinity in the nM range, and the cGi-500 biosensor^21^, which is >10 times less sensitive. The untargeted ScGI and the OMM-ScGI biosensors constructed in this study presented a mid-range affinity with an EC_50_ of around 200 nM, and were shown to be sensitive enough to detect similar increases of cGMP by CNP stimulation compared to the other biosensors (Fig. 2d, Supplementary Fig. 2d). ANP stimulation, however, gave a significant increase in cGMP only at the OMM and not with the untargeted biosensor in cardiomyocytes (Fig. 2d), indicating that GC-A stimulation locally increases cGMP around the OMM. CNP has previously been shown to stimulate GC-B both in the crest and T-tubules and produce cGMP that reaches remote targets in cardiomyocytes^20, 31^, therefore the cGMP increase at the OMM could reflect the widespread CNP-induced cGMP increase. ANP, on the other hand, has been reported to produce a small cGMP pool confined near the plasma membrane by activation of GC-A only in the T-tubules of cardiomyocytes^31^. Such local cGMP increase might not be detected by an untargeted biosensor due to widespread distribution and local signals being “diluted” across the entire cell. In fact, even using the ∼10 times more sensitive untargeted Yellow *Pf*PKG biosensor we could previously only detect a modest cGMP increase after GC-A stimulation^19^. Since ANP increases cGMP considerably more at the OMM nanodomain compared to that detected by the untargeted biosensors in cardiomyocytes (Fig. 2d), the GC-A receptor and therefore its cGMP pool at the T-tubules might be in close vicinity of the OMM and form a nanodomain. Given that mitochondria are tightly distributed between the sarcomeres but also in the sub-sarcolemmal region^35^, either the membranal localization of the GCs or the different ability of cGMP from the GC-A and GC-B to reach different targets could help explain the higher levels of cGMP detected at the OMM by CNP compared to ANP.

### CNB domains responsible for inducing changes in FRET in the cGI biosensors

Given that cAMP is structurally similar to cGMP and that basal cAMP concentration in cardiomyocytes are higher (1µM) than cGMP (10nM)^15–17^, it is important to determine the selectivity of cGMP biosensors and to ensure that the responses observed are representative of increases in cGMP and not cAMP. In our hands, the cGi-500V showed a poor selectivity profile, where cAMP bound to the biosensor at low concentrations resulted in *increased* FRET (Fig. 1a). However, these concentrations of cAMP did not compete with the ability of cGMP to alter FRET (Supplementary Fig. 1a). In addition, when mutating the CNB-A domain (in the ScGI), the increase in FRET by low cAMP concentrations was prevented, whereas the cGMP potency was slightly *increased* (Fig. 1b and Supplementary Fig. 1b). It has previously been shown that the CNB-A domain displays similar high affinity for both cGMP and cAMP^25^, whereas the CNB-B domain has a slightly lower affinity for cGMP (∼215nM), but the affinity for cAMP is much lower (EC_50_=52µM) and the CNB-B domain is therefore responsible for the cGMP selectivity of PKGI^26^. Considering that the ScGI has similar affinity for cGMP as the isolated CNB-B domain and mutating the arginine in β5 in this domain reduced cGMP affinity (Supplementary Fig. 1), we consider this domain as solely responsible for the biosensors sensitivity for cGMP. Taken together, this indicates that 1) the FRET increase seen with low concentrations of cAMP (in the cGi-500V) is possibly due to cAMP binding to the CNB-A domain, 2) cAMP at high concentrations binds to the CNB-B domain of cGi-500V and ScGI, reducing FRET, and 3) cGMP binding to the CNB-A does not alter the conformation of PKG to trigger a change in FRET, whereas cGMP binding to the CNB-B domain triggers an increase in the distance between the CFP and Venus, thus reducing FRET. The ScGI biosensor will thus be a valuable tool for determining cGMP, particularly in complex biological systems where cAMP can act as a confounding factor.

### NPs/cGMP signalling involved in mitochondrial function

Our results show that ANP and CNP increase cGMP at the OMM. Cyclic GMP has previously been found in cardiac mitochondria, where mtNOS and sGC have been reported to be located in the matrix^36–38^ and sGC-generated cGMP has also been found at the OMM^30^. However, those studies did not examine natriuretic peptide signalling.

Here we demonstrate increased survival of adult cardiomyocytes and protection from apoptosis by NP stimulation. Previous studies determining the effects of NPs on regulation of apoptosis show conflicting results: some studies found that stimulating GC-A and/or GC-B is protective against apoptosis, mostly in other cell types^39–42^, but also in cardiomyocytes and the heart^43–45^. However, these latter studies used neonatal cardiomyocytes, which have a lower degree of differentiation, focused on ANP signalling specifically at the nuclei or did not study whether the NP-effect was directly on cardiomyocytes. Others showed an increase in apoptosis by NPs in cardiomyocytes^46–49^ and in other tissues^50^. However, the different results between the studies showing increased apoptosis and the data presented in this study could potentially be due to study of adult (this study) *vs*. neonatal cardiomyocytes^46, 48, 49^ or cardiomyocytes from normal heart (this study) *vs*. a heart failure model^47^.

The exact mechanism of how NPs regulate mitochondrial function and prevent apoptosis remains elusive. Some studies show that NPs promote mitochondrial biogenesis by upregulation of PGC-1α and PPARδ and this mechanism could protect cardiomyocytes against mitochondria-mediated apoptosis^51, 52^. Moreover, PKGI seems to protect against apoptosis by increasing expression of Bcl-2 and decreasing Bax in adult rat cardiomyocytes subjected to ischemia/reoxygenation stress^53^. Further, ANP- and BNP-stimulation has been shown to induce mitoK_ATP_ opening and prevent mitochondrial permeability transition pore opening, where both mechanisms are protective against apoptosis^44, 54–56^. A third pathway for NP regulation of mitochondrial function could be through regulation of Drp1. It has earlier been shown that PKA can phosphorylate Drp1 at S637, reducing its activity ^32^, and this has been associated with prevention of apoptosis ^33, 34^. PKG-mediated phosphorylation of Drp1 and its protective effect has been shown via the NO/cGMP pathway in myogenic precursor cells ^57^. Thus, there are several possible pathways for how natriuretic peptides can regulate apoptosis, involving either gene regulation or direct regulation of apoptotic pathways. In the present study, we show that NPs increase phosphorylation of Drp1 at Ser637. This could prevent Drp1 activation and initiation of mitochondrial fragmentation, as shown by CNP stimulation (Fig. 3b), and thus possibly cause prevention of apoptosis^58^.

There are some indications for the presence and effect of cGMP inside the mitochondrial matrix and at the OMM after stimulation of sGC^30, 36–38, 44, 54–56^. However, the role of NP-stimulated cGMP at the OMM shown herein has not been examined previously. Although both CNP and ANP showed comparable effects on LDH activity in the medium, ATP production, PARP cleavage, cleaved caspase 9 and cytochrome c release in addition to the prevention of apoptosis (Fig. 4-5), stimulation of CNP gave both higher cGMP levels at the OMM and concomitantly larger phosphorylation of Drp1 compared to ANP-stimulation (Fig. 2 and 3), in addition to reduced H_2_O_2_-induced mitochondrial fragmentation (Fig. 3). This might indicate that either different pathways are involved in the anti-apoptotic effects seen after ANP and CNP stimulation, or that other pathways are also involved. Alternatively, the magnitude of Drp1 phosphorylation seen after ANP stimulation is sufficient to inhibit the mitochondrial intrinsic pathway.

In conclusion, we constructed a novel FRET-based biosensor with high cGMP selectivity in order to detect cGMP at the mitochondria and show that both CNP and ANP increase cGMP at the OMM. Further, we present evidence for a protective effect of natriuretic peptides against cardiomyocyte apoptosis, mediated by inhibition of the intrinsic apoptotic pathway, which also involves phosphorylation of Drp1 and reduced mitochondrial fragmentation.

## Methods

### Materials

Human C-type natriuretic peptide (CNP) and rat atrial natriuretic peptide (ANP) were from GenScript Corp. (Piscataway, NJ, USA). Mitotracker Deep Red, Mitotracker Green, Dulbecco’s modified Eagle’s medium (DMEM) with GlutaMAX, sodium pyruvate and MEM NEAA were from ThermoFisher Scientific. All other chemicals and media were from Sigma-Aldrich (Sigma-Aldrich, St.Louis, MO), unless otherwise specified.

### Animals

This investigation was approved by the Norwegian Food Safety Authority (application number 8683 and 23286) and conforms to the research animal Directive 2010/63/EU, the Guide for the Care and Use of Laboratory Animals (NIH publication No. 85-23, revised 2011, US), in accordance with the ARRIVE guidelines^59^. Animals were anaesthetized in 4-5 % isoflurane and anesthesia was confirmed by abolished pain reflexes. Animals were euthanized by cervical dislocation and the chest was then opened to extract the heart.

### Isolation of cardiomyocytes

Primary culture of cardiac ventricular myocytes from male Wistar rats were prepared as previously described^60^.

### LDH measurements

Cell death was measured by the LDH-Glo™ Cytotoxicity Assay (Promega, Mannheim, Germany). Cardiomyocytes cultured for 20 h were stimulated with CNP (1 µM), ANP (1 µM) or vehicle (H_2_O) at time 0. Medium was sampled at the indicated time (0, 3, 6, 12 and 24 h) and added to LDH storage buffer (200 mM Tris-HCl pH 7.4, 10% glycerol, 1% BSA). Measurements were performed according to manufacturer’s instructions.

### ATP assay

Cardiomyocyte viability was measured as changes in cellular ATP levels using the CellTiter-Glo® 2.0 Cell Viability Assay (Promega). Cardiomyocytes cultured for 20 h were stimulated with CNP (1 µM), ANP (1 µM) or vehicle (H_2_O). After 3 and 24 h from stimulation, cardiomyocytes were washed with 50 mM Tris-HCl pH 7.4, 150 mM NaCl, 2 mM EDTA and lysed using Solobuffer cell lysis kit (Fabgennix, Frisco, TX 75033, US). Measurements of ATP in the samples were performed according to manufacturer’s instructions. Change in cellular ATP content after each stimulation time was calculated as (ATP in treated cells)/(ATP in vehicle-treated).

### TUNEL assay

After 20 h from plating, adult cardiomyocytes were stimulated with either CNP (1 µM), ANP (1 µM) or vehicle (H_2_O) for 3 h. To detect apoptotic cells, the ApopTag Plus Fluorescein In Situ Apoptosis Detection Kit (Merck, Darmstadt, Germany) was used according to the manufacturer’s protocol. DAPI (5µg/ml) was added for nuclear staining. Cells were visualized under an Olympus BX53 (Olympus, Japan), using an air objective (10x, 0.30NA, Olympus), and images taken with a XC50 color camera and cellSens Entry Software (Olympus, Japan). Images were acquired and analysed by a blinded operator.

### Cytoplasmic cytochrome c release

Adult cardiomyocytes were cultured for 20 h and then treated with CNP (1 µM), ANP (1 µM) or vehicle (H_2_O) for 3 h, washed twice with 50 mM Tris-HCl pH 7.4, 150 mM NaCl, 2 mM EDTA, incubated on ice with “cytoplasmic extraction buffer” from Subcellular Protein Fractionation Kit (Thermo Scientific, Waltham, USA) for 15 min, cells were then scraped and samples collected and centrifuged at 4 °C for 5 min at 500 × g. The supernatant (cytoplasmic fraction) was further centrifuged at 13000 × g for 10 min to separate the mitochondria (and other organelles). The supernatant was then collected and cytochrome c measured using cytochrome c ELISA Kit (MyoBiosource, San Diego, USA) following the manufacturer’s instructions.

### Immunoblot analysis

Cardiomyocytes cultured for 20 h were stimulated with CNP (1 µM), ANP (1 µM) or vehicle (H_2_O). After 3 h stimulation, protein was extracted on ice using lysis buffer (Tris 50 mM, EDTA 1 mM, EGTA 1 mM, NaF 50 mM, DTT 1 mM, Na_4_P_2_O_7_ 10 mM, Na_3_VO_4_ 1 mM, Triton 100 0.01%, SDS 0.05% and deoxycholat 0.5 %) supplemented with protease inhibitor cocktail and PMSF (1 mM). Samples were solubilized in loading buffer (62.5 mM Tris-HCl, glycerol 10%, bromophenol blue 1% wt/vol, SDS 2%, mercaptoethanol 2.5%) and denatured for 5 min at 95°C. Protein samples were separated by SDS-PAGE using 6%, 7.5% or 15% polyacrylamide gels and then transferred to a PVDF membrane. Membranes were blocked with 3% non-fat dry milk in Tris-buffered saline containing Tween-20 (0.1%) for 2 h and then incubated with the indicated primary antibodies overnight at 4°C, followed by incubation with appropriate secondary antibodies for 1 h at room temperature. Drp1 expression was quantified using a nitrocellulose membrane previously probed with pDrp1 S637. The membrane was stripped for 10 min in stripping solution (Glycine 0.15% w/v, SDS 0.1% w/v, Tween-20 1%, pH 2.2), then washed with phosphate-buffered saline and Tris-buffered saline containing Tween-20 (0.1%). The membrane was then blocked and incubated with Drp1 primary antibody followed by appropriate secondary antibody as described above. Blots were developed by enhanced chemiluminescence reaction using an Epi Chemi II Darkroom with Labworks Software (UVP Laboratory Products, Cambridge, UK). Protein expression or protein phosphorylation was quantified as measured intensity of area under the curve in ImageJ software. The intensity of each band was normalized to the intensity of the corresponding loading control (vinculin) except for cleaved PARP that was normalised to total PARP (cleaved+uncleaved). Antibodies: Vinculin (1:5000), PARP (1:1000), Cleaved PARP (1:800), Caspase 9 (1:1000), Phospho Drp1 S637 (1:1000), Drp1 (1:1000), all from Cell Signaling.

### Construction of FRET-based biosensors for cGMP

The sequences encoding cGi-500V and ScGI sensors were synthesized by GenScript and inserted into the vector pcDNA3.1(-). cGi-500V sequence was from *Bos Taurus* PKG I (Gln^79^-Tyr^345^) flanked by CFP and Venus. The ScGI sequence differs only by the mutation Cys158Arg. The OMM-targeting sequence AKAP1 (1-30), was synthetized by GenScript, opened by *Eco*RI*/Xba*I and inserted N-terminally into the ScGI sequence preceded by a linker A(EAAAK)^5^A to separate the fusion proteins and keep the flexibility^61^. The nucleotide sequences for ScGI has been deposited in GenBank under the accession number OP221137. ScGI and OMM-ScGI biosensors were then packaged in adenovirus type 5, amplified and titer measured by VectorBiolabs (Malvern, PA).

### Transfection of HEK293 cells and *in vitro* FRET assay

HEK293 cells were cultured in Dulbecco’s modified Eagle’s medium (DMEM) with GlutaMAX supplemented with 10% fetal bovine serum (FBS) and 100 U penicillin/streptomycin at 37°C in a humidified atmosphere of 5% CO_2_ in air. Cells were transiently transfected with the indicated plasmids using LipofectAMINE 2000 (ThermoFischer) according to the manufacturer’s protocol. After transfection, cells were cultured for 48 h, then homogenized with Ultra-Turrax (Franke & Kunkel KG, Staufen, Germany) for 30 s at 4°C in Buffer A (HEPES 10 mM pH 7.3, NaCl 137 mM, KCl 5.4 mM, CaCl_2_ 2 mM, MgCl_2_ 1 mM). Homogenates were stimulated with increasing concentrations of cGMP or/and cAMP in the presence of IBMX (100 µM). The donor fluorophore CFP was excited at 430 nm (430/24 filter) and the fluorescence emission (F) of Venus and CFP were measured (535/25 and 470/24 nm, respectively) in an EnVision plate reader (PerkinElmer, USA, NY, Melville). Venus emission was corrected for spillover of CFP emission into the Venus channel (72.3%). The FRET response was calculated as changes in F_Venus_/F_CFP_. Affinities (*p*EC_50_) were calculated by GraphPad Prism 9 for Windows (GraphPad Software, San Diego, CA) where the lowest FRET was set as the highest concentration of cGMP.

### Confocal microscopy

#### Evaluation of the FRET biosensors localization at mitochondria

Rat heart derived H9c2 cardiac myoblasts (Sigma-Aldrich, St.Louis, MO) were cultured in Dulbecco’s modified Eagle’s medium (DMEM) with GlutaMAX supplemented with 10% fetal bovine serum (FBS), 100 U penicillin/streptomycin, 100 mM sodium pyruvate and 100 mM MEM NEAA at 37°C in a humidified 5% CO_2_–95% air atmosphere. Cells were transiently transfected with the indicated plasmids using LipofectAMINE 2000 (ThermoFisher) according to the manufacturer’s protocol and cultured on poly-lysin coated glass cover-slides. Rat adult cardiomyocytes were plated in laminin coated cover-slides and transduced with adenovirus containing the biosensor as previously described^62, 63^. After 48 h transfection (H9c2) and transduction (cardiomyocytes), cells were incubated with 50 nM Mitotracker Deep Red (ThermoFisher) for 30 min, washed and incubated in Buffer B (NaCl 144 mM, KCl 1.97 mM, CaCl_2_ 1 mM, MgCl_2_ 1 mM, KH_2_PO_4_ 0.43 mM, K_2_HPO_4_ 1.5 mM, glucose 10 mM) to be visualized under an Olympus FV1000/BX61 confocal microscope using a water immersion objective (60x 1.1 NA). Cells were sequentially excited with a 488 nm laser for CFP and a 633 nm laser for the Mitotracker Deep Red and the emission was measured by Olympus Fluoview (Olympus, Tokyo, Japan) at 510/25 nm for CFP and 664/20 nm for Mitotracker Deep Red.

#### Evaluation of mitochondria morphology

H9c2 cells were cultured on poly-lysin coated glass cover-slides and after 24 h stimulated as indicated and incubated with 50 nM Mitotracker Green for 15 min, washed and incubated in Buffer B. Cells were visualized under an Olympus FV1000/BX61 confocal microscope using a water immersion objective (60x 1.1 NA). Cells were excited with a 488 nm laser and the emission was measured by Olympus Fluoview 1000 at 510/25 nm. Images were analysed by Mito-Morphology ImageJ macro^64^, which was modified for this measurements to apply a Gaussian filter. In brief, the macro inverts the green channel (Mitotracker Green) to a binary mask and thresholds to resolve mitochondria; the size and shape are analysed as particles and the ratio between the major and minor axis for each mitochondria was employed as an index of mitochondria elongation. For each cell the median of this mitochondrial elongation was calculated and presented.

### FRET imaging

H9c2 cells and cardiomyocytes expressing the indicated biosensors (as described above) were placed into a watertight cell imaging chamber (Attofluor, ThermoFisher) at room temperature in Buffer B. Cells were visualized through a motorized digital inverted fluorescent microscope (iMIC; FEI, Graefelfing, Germany) with an oil objective (40x, 1.35 NA; Olympus). Fluorescent molecules were illuminated by a monochromator with fast-switching wavelengths (Polychrome V; FEI) for 20-80 ms at a frequency of 0.1-1 Hz. Cells were excited at 436±10 nm for 20-80 ms and emission was split using a dichrotome iMIC Dual Emission Module with a DCLP 505 filter that separates the two fluorophores onto a camera (EVOLVE 512; Photometrics, Tucson, USA). Fluorescence was measured at 530±15 nm for Venus and at 470±12 nm for CFP emission. Images were acquired by Live Acquisition browser (FEI) and FRET was calculated using Offline Analysis software (FEI). FRET was measured as F_Venus_/F_CFP_. Venus emission was corrected for spillover of CFP emission into the Venus channel (63%). FRET was adjusted for photobleaching. In a subset of experiments, cells were visualized through a motorized digital inverted microscope (IX81; Olympus) with an oil objective (40x, 1.35 NA; Olympus). Fluorescent molecules were excited at 436nm for 8-20ms (CoolLed pE-4000) and emission split with a 495 nm filter using an Optosplit II (Cairn research, Kent, UK) separating two fluorophores onto a camera (Prime 95B; Photometrics). Fluorescence was measured at 535±15 nm for Venus and at 470±12 nm for CFP emission. Images were acquired by Micro-Manager. FRET was measured as F_Venus_/F_CFP_ through custom-built scripts in ImageJ. Representative experiments were run through a Savitzky–Golay filter using GraphPad Prism 9 for Windows for clarity.

### Total cGMP measurements

H9c2 cells and isolated ventricular cardiomyocytes plated for 20h were stimulated with ANP (1µM) or CNP (1µM) for the indicated time. Cyclic GMP levels were measured using a cGMP enzyme immunoassay kit (Cayman Chemical Company, Ann Arbor, MI, USA) as previously described ^65^.

### Statistics

Statistical significance was determined by one-way ANOVA with Bonferroni’s, Dunett’s, Tukey’s and Sidak’s multiple comparisons test, two-way ANOVA with Sidak’s multiple comparisons test as indicated using GraphPad Prism 9 for Windows. P<0.05 was considered statistically significant.

### Data availability

The nucleotide sequences for ScGI has been deposited in GenBank under the accession number OP221137. The authors declare that the data supporting the findings of this study are available within the paper and its supplementary information files.

## Acknowledgements

The authors wish to thank Iwona Gutowska Schiander (University of Oslo) for excellent technical assistance and Soheil Naderi (Oslo University Hospital) for valuable advice, help and discussions. This work was supported by the South-Eastern Norway Regional Health Authority (grant # 2015036, 2019051 and 2023045), the Norwegian Council on Cardiovascular Diseases, The Research Council of Norway (grant # 205167 and 303490), The Anders Jahre Foundation for the Promotion of Science, The Family Blix Foundation, The Simon Fougner Hartmann Family Foundation, and grants from the University of Oslo

## Author Contribution

GC, BDN, CK, KWA and LRM designed research; GC, BDN, DA and MO performed research. GC, BDN, DA, MO, FOL, KWA and LRM analyzed data; GC, FOL, KWA and LRM wrote the manuscript. All authors revised and approved the manuscript.

## Supplementary Information

Supplementary Tables and Figures.

