## Supplementary Tables and Figures for "Natriuretic peptides increase cGMP around cardiomyocyte mitochondria and protect against apoptosis"

##### Supplementary Table

##### Supplementary Table 1

**Selectivity of the FRET biosensors.** HEK293 cells expressing the indicated biosensor were homogenized and incubated with increasing concentration of cGMP with or without cAMP at the indicated concentrations and fluorescence measured as described in Materials and Methods. $EC_{50}$ values shown in µM were calculated from the average $pEC_{50}$ values. Results shown are mean±SEM of 5 experiments. *p<0.001 *vs*. cGi-500V; ‡p<0.01 for cGMP $EC_{50}$ *vs*. cGMP $EC_{50}$ +100µM cAMP (One-way ANOVA, with Sidak's post hoc test).

| | **cGMP** | | **cGMP +1µM cAMP** | | **cGMP +100µM cAMP** | |
|---|---|---|---|---|---|---|
| **Sensor** | $pEC_{50}$ | $EC_{50}$ (µM) | $pEC_{50}$ | $EC_{50}$ (µM) | $pEC_{50}$ | $EC_{50}$ (µM) |
| cGi-500V | 6.33±0.05 | 0.5 | 6.52±0.08 | 0.3 | 5.94±0.06‡ | 1.1 |
| ScGI | 6.7±0.05* | 0.2 | 6.65±0.16 | 0.2 | 6.24±0.14‡ | 0.6 |

### Supplementary Figures

#### Supplementary Figure 1

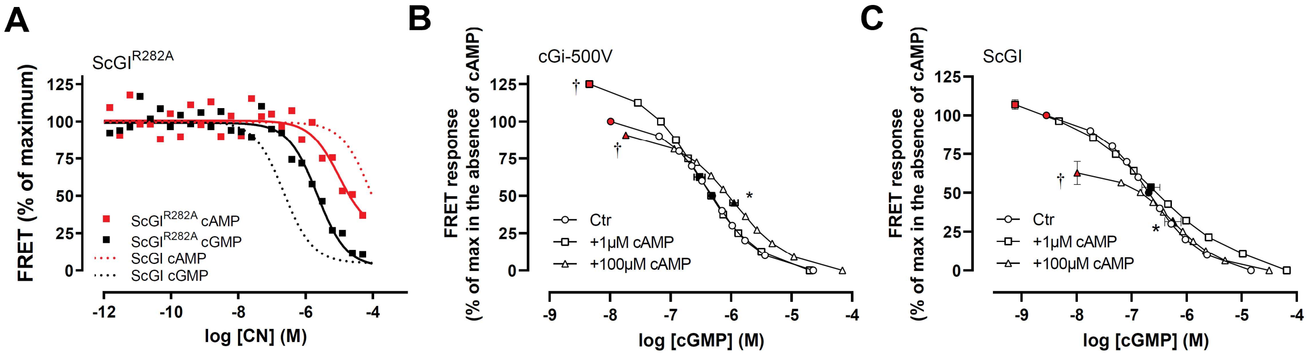


**Supplementary Figure 1. CNB-B domain is responsible for the cGMP-induced FRET change of the ScGI biosensor and the CNB-A domain is responsible for the high cAMP affinity of the cGi-500V. A)** Homogenates of HEK293 cells transfected with the mutant $ScGI^{R282A}$ biosensor were incubated with increasing concentrations of either cGMP or cAMP. FRET ($F_{Venus}/F_{CFP}$) was measured as described in Materials and Methods and normalized to FRET in the absence of cGMP and at the highest cGMP concentration. Shown are representative of 4 independent experiments. The affinities ($pEC_{50}$) of cGMP and cAMP were 5.68 ± 0.05 and 5.02 ±0.06, respectively, and the maximal change in FRET was 19,6 ±1,2 % from 4 independent experiments. **B)** and **C)** Concentration-response curves generated by increasing concentrations of cGMP in homogenates of HEK293 cells expressing either the cGi-500V **(B)** or the ScGI **(C)** biosensor in the absence (Ctr) or presence of the indicated concentration of cAMP. Affinity for cGMP ($pEC_{50}$; shown as black filled symbols) and FRET in the absence of cGMP (shown as red filled symbols) are shown as mean±SEM from 4-5 independent experiments *p<0.01 for $EC_{50}$ of +100µM cAMP *vs*. Ctr (one-way ANOVA, Sidak's test). †p<0.05 for % change in FRET of indicated condition *vs*. Ctr (one-way ANOVA, Dunnett's multiple comparison test).

#### Supplementary Figure 2

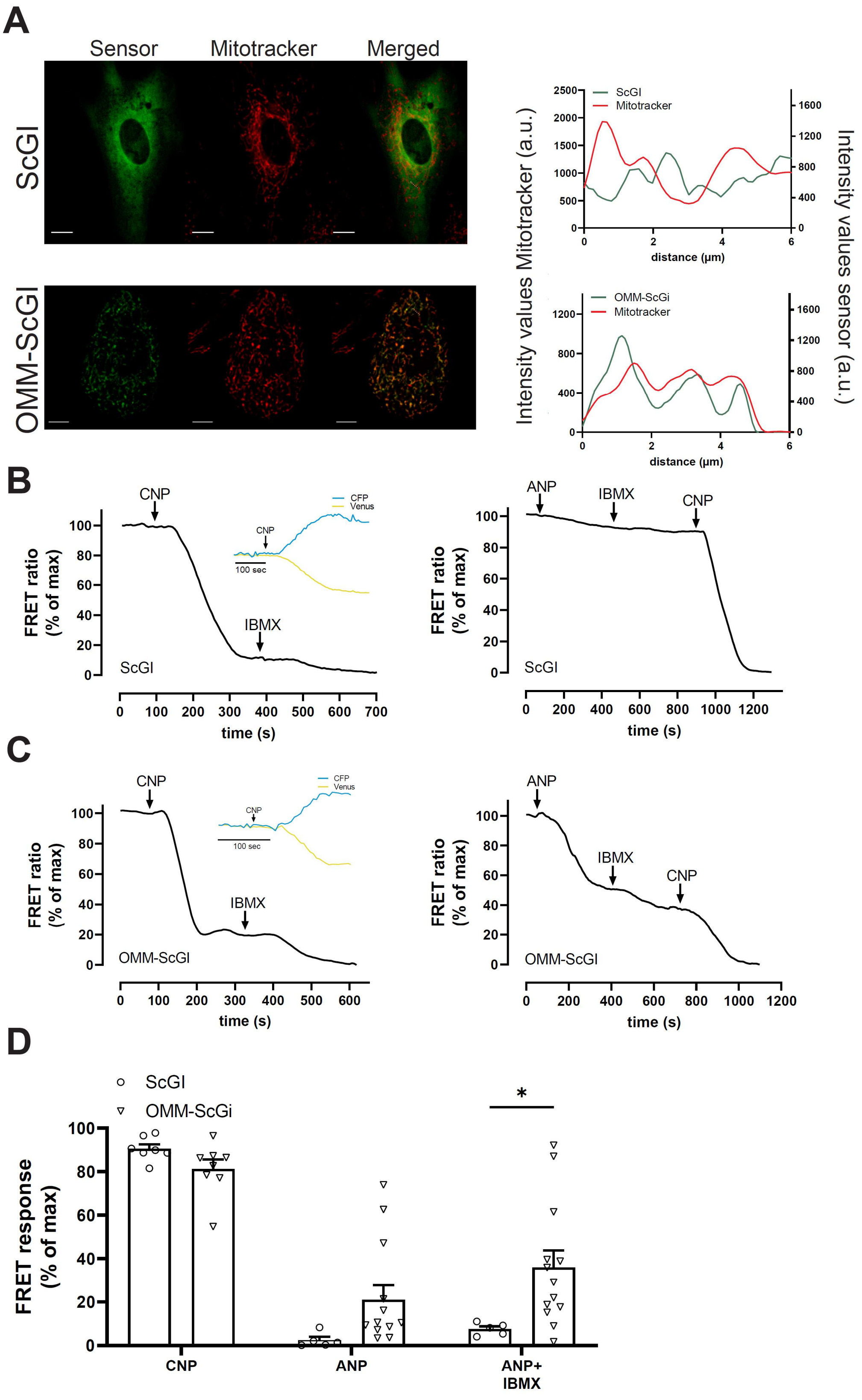

**Supplementary Figure 2: CNP and ANP increase cGMP at the outer mitochondrial membrane of H9c2 cells. A)** Cells expressing the ScGI (upper panel) and OMM-ScGI (lower panel), were stained with Mitotracker Deep Red, washed and visualized under a confocal microscope. Scale bar: 10µm. Right panels show colocalization by intensity overlay of both fluorescent signals (Mitotracker and sensors) from indicated dotted line. **B and C)** Left panels show representative of FRET ratio (Venus/CFP) in single cells expressing the indicated biosensor and stimulated with CNP (300nM) and IBMX (100µM) (left panels), or ANP (1µM) followed by maximal activation of the biosensor (IBMX 100µM + CNP 300 nM) (right panels). The inserts in the left panels show corresponding traces of separate Venus and CFP intensities upon stimulation with CNP (300 nM). Shown are traces representative of 5-14 cells. **D)** Quantification of FRET responses from CNP-, ANP- and IBMX + ANP- stimulation. *;p<0.01 (one-way ANOVA, Sidak test).

#### Supplementary Figure 3

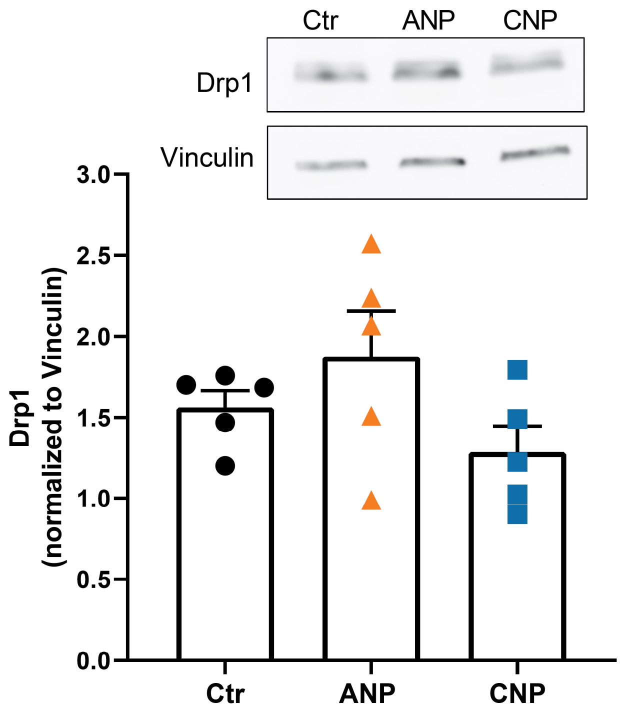


**Supplementary Figure 3. CNP and ANP do not change Drp1 expression levels.** Adult cardiomyocytes were stimulated with Vehicle (Ctr), CNP (1 µM) or ANP (1 µM) for 3 h and lysed and Drp1 expression levels were determined as described in Materials and Methods and normalized to loading control (vinculin). Insert: representative western blots. Lower panel: quantification of blots presented as mean±SEM from 5 independent experiments. Stimulation with ANP or CNP were not significantly different from Ctr ($p=0.37$ and $p=0.16$, respectively).

#### Supplementary Figure 4

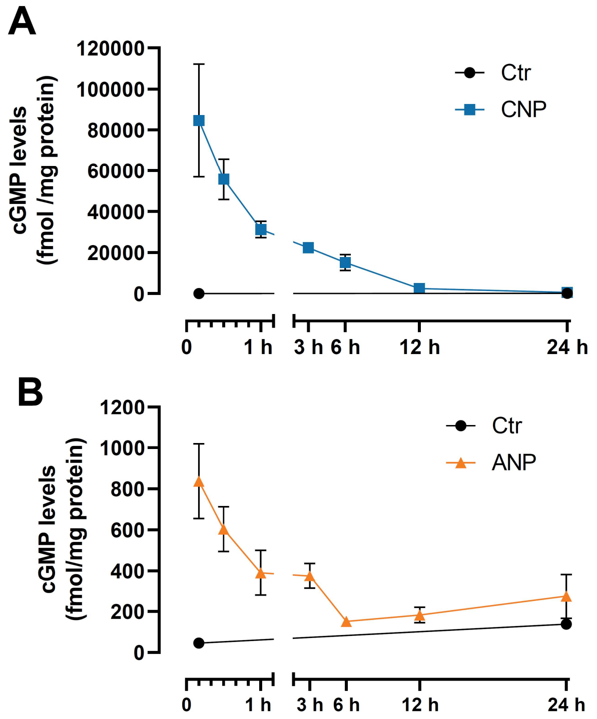


**Supplementary Figure 4: CNP and ANP increase cGMP over 12 h in H9c2 cells.** Total cGMP levels in H9c2 cells in the absence (Ctr) or presence of **A)** CNP (1µM) or **B)** ANP (1µM), at the indicated stimulation times. Data are mean±SEM from 3 individual experiments.

#### Supplementary Figure 5

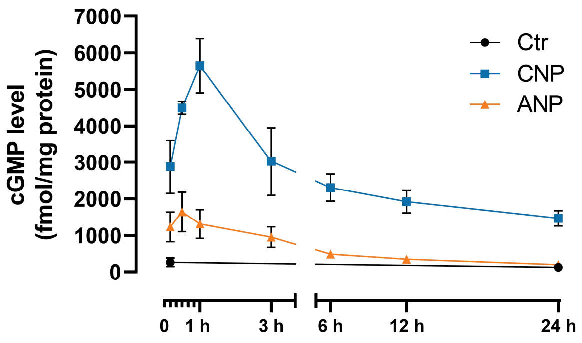


**Supplementary Figure 5. CNP and ANP increase cGMP in cardiomyocytes.** Total cGMP levels in isolated cardiomyocytes in the absence (Ctr) or presence of either ANP (1µM) or CNP (1µM), at the indicated stimulation times. Data are mean±SEM from 3 individual experiments.
